# Inhalation of nanoparticles during pregnancy enhances placental glucose transport in rats

**DOI:** 10.64898/2026.06.16.732724

**Authors:** Talia N. Seymore, Sara Hoffmann, Pedro Louro, Carol Gardner, Michael J. Goedken, Phoebe A. Stapleton

**Author notes:** **Corresponding author** Phoebe Stapleton, PhD, ATC, Environmental and Occupational Health Sciences Institute (EOHSI), Ernest Mario School of Pharmacy, Rutgers University, 170 Frelinghuysen Rd., Piscataway, NJ 08854.

## Abstract

Fetal health is heavily dictated by the maternal environment. Inhaling airborne pollutants, like particulate matter, is associated with pregnancy complications and fetal developmental pathologies, including fetal growth restriction (FGR). Because fetal growth is dependent on the placental transfer of nutrients from the maternal circulation, particularly glucose, investigating glucose transport capacity is critical to understanding the development of FGR associated with gestational inhalation of particulate matter. Pregnant Sprague Dawley rats were exposed to titanium dioxide nanoparticles (9.8±1.0 mg/m^3^) as a proxy for ultrafine particulate matter, from gestational day (GD) 5 to GD 19 via whole-body inhalation. Glucose transporters (GLUTs) 1, 3 and 4 were evaluated in term placentas on GD 20 and *ex vivo* placental perfusion was conducted as a functional assessment of glucose transport. Exposure resulted in a reduction in *Glut3* mRNA and GLUT1 protein. However, exposed placentas exhibited an adaptation, characterized by increased GLUT4 expression and membrane localization of both GLUT1 and GLUT4. Placental perfusion confirmed these molecular changes, revealing increased glucose flux in exposed placentas compared to control (AUC 95% CI: 77.4 to 127.5 vs 39.1 to 73.6, respectively). Contrary to our hypothesis, exposure to these nanoparticles enhanced glucose transport across the placenta. Here we have demonstrated that inhaling airborne pollutants during pregnancy modulates placental function and nutrient transport mechanisms, which can have direct effects on fetal development. Furthermore, we provide evidence for targeted interventions, aimed at mitigating fetal developmental pathologies.

**Highlights:**

- Gestational inhalation of nanoparticles decreases GLUT1 expression in the placenta.
- The placenta adapts to gestational nanoparticle inhalation by enhancing GLUT4 expression and GLUT1 and GLUT4 membrane localization.
- *Ex vivo* placental perfusion demonstrated increased glucose flux across to the placenta to the fetus following gestational inhalation of nanoparticles.

## Introduction

Fetal development is sensitive to maternal health and the uptake of toxicants from maternal surroundings and is therefore heavily influenced by the maternal environment (Ferguson et al. 2014; Gui et al. 2022; Park et al. 2023; Rokoff et al. 2018; Veras et al. 2009; Vigeh et al. 2023). In humans, exposure to fine and ultrafine particulate matter within air pollution is associated with fetal growth restriction (FGR), and other fetal developmental pathologies as well as pregnancy complications (Dadvand et al. 2013; Gustavsson et al. 2025; Jedrychowski et al. 2010; Niu et al. 2022). FGR occurs when the developing fetus fails to achieve its optimal growth. This outcome is associated with increased risk of perinatal morbidity, mortality, and long-term health effects including cardiovascular, metabolic, and neurological diseases or disorders (Barker 2006; D’Agostin et al. 2023; Vayssière et al. 2015). Several factors can contribute to FGR including maternal undernutrition (Belkacemi et al. 2010) and impaired placental function (Dumolt et al. 2021).

The placenta is a fetal-derived organ developed during pregnancy to regulate nutrient and waste exchange between the mother and fetus (Jarzembowski 2014). Much of this nutrient transport takes place in the labyrinth zone of the rodent placenta (Furukawa et al. 2018). It is highly vascularized and its transport capacity relies on the establishment of an efficient uteroplacental circulation (Wang 2010). Current literature supports an association between maternal inhalation of air pollution contaminants (e.g. ozone, particulate matter) and reduced uteroplacental blood flow in rodents (Cary et al. 2023; Garner et al. 2022; Griffith et al. 2023; Stapleton et al. 2013). Collectively, these findings suggest that gestational exposure to airborne pollutants may impair placental perfusion, thus restricting nutrient bioavailability, and placental nutrient transfer.

As fetal growth relies heavily on glucose metabolism (Battaglia and Meschia 1978), examining glucose transport mechanisms in the placenta is critical to understanding particulate matter- induced FGR. Cellular uptake and transport of glucose occur through glucose transporters (GLUTs). In term placentas from both rats and humans, GLUT1 is the primary transporter for glucose and is highly localized to membranes of the syncytiotrophoblasts (Ericsson et al. 2005; Shin et al. 1997). GLUT3 is less abundant and most important in early pregnancy but decreases in expression throughout gestation (Brown et al. 2011; Shin et al. 1997). Although not very abundant and mostly intracellular, GLUT4 is also present in the placenta and responds to both maternal and fetal insulin stimulation with translocation to syncytiotrophoblast membranes for glucose uptake (James-Allan et al. 2019). Alterations in placental GLUT expression have been associated with a number of environmental pollutants including cadmium, bisphenol A, and air pollution (Ganguly et al. 2024; Molangiri et al. 2024; Yue et al. 2025; Zhu et al. 2021). However, it is unclear at this time if these adaptations associated with particulate matter exposure alter placental function, and more importantly, nutrient transport.

As a fetal-derived organ, the placenta is sexually dimorphic and can have sex-related responses to environmental stressors, including modifications in glucose uptake (Clifton 2010; Gonzalez et al. 2018; Sun et al. 2020; Walker et al. 2014). The purpose of this study was to determine the sex-specific impact of maternal inhalation of aerosolized particulates on placental glucose transport. Pregnant Sprague Dawley rats were exposed via whole-body inhalation to titanium dioxide nanoparticles (nano-TiO_2_) as a surrogate for ultrafine particulate matter. Glucose transporters were assessed in term placentas and real-time transport was evaluated using *ex vivo* placental perfusion. We hypothesized that exposure to aerosolized nanoparticles during gestation reduces glucose transport across the placenta, reducing fetal glucose bioavailability, and promoting the development of FGR.

## Methods

### Animals and Exposure

Fifty-two Pregnant Sprague Dawley rats were purchased from Charles River Laboratories (Kingston, NY) and delivered on gestational day (GD) 4 to an AAALAC accredited vivarium and food and water were provided *ad libitum*. All protocols were approved by Rutgers IACUC. After 24 hr acclimatization, rats (n=26 dams) were exposed to an average concentration of 9.8±1.0 mg/m^3^ of aerosolized nano-TiO_2_ powder (Aeroxide TiO_2_, Parsippany, NJ) for 4-5 hr/d, 5d/wk from GD 5 to GD 19 using a custom designed 84 L whole-body inhalation exposure chamber (IEStechno, Morgantown, WV) as previously described (Fournier et al. 2019; Nurkiewicz et al. 2008; Stapleton et al. 2013; Stapleton et al. 2015b; Yi et al. 2013). Particle size was measured by electrical mobility using a Scanning Mobility Particle Sizer (SMPS, TSI, Shoreview, MN), which captured particles 1-1000 nm in diameter. Average particle size was calculated as 118 ± 1.7 nm (Figure 1A). A High Resolution Electrical Low-Pressure Impactor (HR-ELPI, Dekati, Finland) using aerodynamic diameter and optical counting indicating a peak mode particle size of 188.9 nm (Figure 1B). A target concentration of 10 mg/m^3^ was utilized to represent maximum permissible exposure limits for occupational airborne chemical contaminants (Title 8, Article 107) (Inc. 2024). To identify if the airflows within the inhalation facility alone impacted placental development, a separate group of control animals were placed into the facility and exposed to filtered air (n=4). We identified no significant differences GLUT protein expression (Supplemental Figure 1) between “filtered air” and naïve control animals that did not enter the inhalation facility. These data are in line with previous findings identifying no significant differences in uterine artery vascular function between “filtered air” and naïve groups (Fournier et al. 2019). Therefore, the “filtered air” results (n=4 dams) were combined with the naïve outcomes (n=22 dams) to form the control group (n=26 dams) presented within this study, as previously described (Fournier et al. 2019).

**Figure 1.**
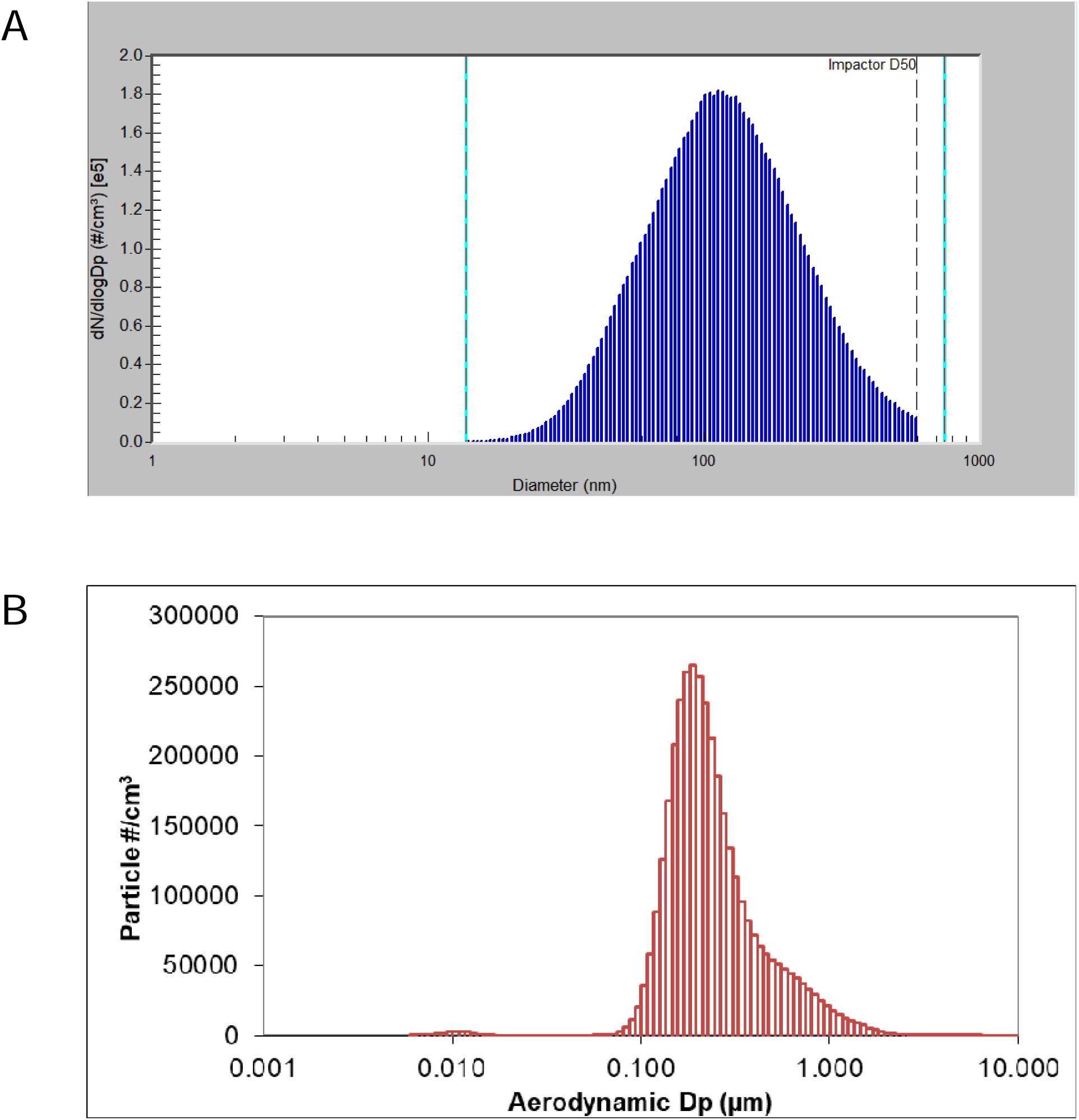
Real-time particle size characterization using electrical mobility (A; SMPS) and aerodynamic diameter (B; ELPI).

### Tissue Collection

After an overnight fast (16 hr), rats were anesthetized with isoflurane (5% induction, 3% maintenance) and weighed on the morning of GD 20. Blood glucose was measured from the tail vein using a commercially available glucometer (One Touch Ultra 2). Mean arterial blood pressure (MAP) was measured using a BLPR2 pressure transducer (World Precision Instruments, Sarasota, FL) and blood was collected via direct carotid cannulation into EDTA-containing vacutainers (Becton Dickinson) and centrifuged at 1100 xg for 10 min. Dams were then euthanized via pneumothorax and heart removal. Plasma from a subset of animals (n=8 dams) was snap frozen in liquid nitrogen and stored at −80 °C for future analyses. Fetal and placental weights were recorded, and placental efficiency was calculated by dividing fetal weight by placental weight (FW/PW) (n=15 litters). Fetal blood glucose was measured via glucometer as described above in trunk blood after decapitation (n=16 litters). Placentas from randomized intrauterine positions on the right and left horns were collected (n=11 litters) and snap frozen in liquid nitrogen and stored at −80°C until RNA isolation or fixed in 10% neutral buffered formalin for histological examination. Maternal insulin concentrations were evaluated (n=8 dams) using a commercially available ELISA kit (Thermo Fisher ERINS).

### RNA Extraction and Real-Time Quantitative PCR

Placental RNA was extracted from two male and two female placentas per dam (n=7 litters) using TRIzol Reagent (Invitrogen^TM^ 15596018) and chloroform. RNA integrity was evaluated using a NanoDrop and confirmed by visualization of 28S and 18S rRNA bands on denaturing gels. Complimentary DNA (cDNA) was generated from RNA (13.2 µg) using Applied Biosystems^TM^ High-Capacity cDNA Reverse Transcription Kit and Power SYBR^TM^ Green PCR Master Mix used to measure amplified signals of *Glut1* (*Slc2a1*) (NCBI Reference Sequence: NM_138827.2), *Glut3* (*Slc2a3*) (NCBI Reference Sequence: NM_017102.3), and *Glut4* (*Slc2a4*) (NCBI Reference Sequence: NM_012751.2). Amplification cycling was conducted using QuantStudio Applied Biosystems ViiA7 qPCR machine and *Glut* CT values were normalized to *B-actin* (NCBI Reference Sequence: NM_031144.3). Fold changes were calculated relative to the average of female naïve samples. Primer sequences are listed below:

*Glut1* forward: 5’-ACG TCC ATT CTC CGT TTC AC-3’

*Glut1* reverse: 5’-TCC CAC GGC CAA CAT AAG-3’

*Glut3* forward: 5’-GAC CAA GCG ACG GAG ATC-3’

*Glut3* reverse: 5’-AGA GCT CCA GCA CAG TGA CC-3’

*Glut4* forward: 5’-AGG CAC CCT CAC TAC CCT TT-3’

*Glut4* reverse: 5’-ATA GCC CTT TTC CTT CCC AA-3’

*B-actin* forward: 5’-ATG TAC CCA GGC ATT GCT GA-3’

*B-actin* reverse: 5’-AGG GTG TAA AAC GCA GCT CA-3’

### Immunostaining

All immunohistology samples were formalin fixed, paraffin embedded, sectioned, hematoxylin and eosin stained, and reviewed by a third-party, board-certified veterinary pathologist for overt pathologies.

Placental tissue from two male and two female fetuses per dam (n=11 litters) were fixed in 10 % neutral buffered formalin and then paraffin embedded. Five µm sections were prepared and deparaffinized, dehydrated through a graded ethanol series, and subjected to heat-induced antigen retrieval with citrate buffer (pH 6.0) for 20 min in a steamer. Sections were then incubated with rabbit monoclonal anti-GLUT1 (ab115730, 1:2500, Abcam, Cambridge, MA), rabbit monoclonal anti-GLUT3 (MA5-32697, 1:1500, Invitrogen, Frederick, MD), or rabbit polyclonal anti-GLUT4 (PA5-80022, 1:200, Invitrogen), after which tissue sections were incubated with secondary antibody horse anti-rabbit IgG polymer (Vector Laboratories, Burlingame, CA, MP6401) for 30 min. Antibody binding was visualized using diaminobenzidine (DAB) chromogen substrate (SK-4100, Vector Laboratories) followed by one min incubation in hematoxylin. Images of tissue sections were acquired at 40x using a Zeiss Axioscan 7. DAB intensity was quantified from two locations (430 µm by 430 µm each) in the labyrinth zone of each tissue using ImageJ Fiji Software (Crowe and Yue 2019).

For immunofluorescence, slides were heated to 60 °C, deparaffinized, and rehydrated through a graded ethanol series, and subjected to heat-induced antigen retrieval with Tris-EDTA buffer (10 mM tris base, 1 mM EDTA, pH 9.0) for 10 min in a steamer. Sections were then incubated with Alexa Fluor 488 wheat germ agglutinin (WGA) (W11261, 5 µg/mL, ThermoFisher) for 10 min to visualize cell membranes for GLUT localization, then incubated with rabbit monoclonal anti-GLUT1 (1:500, Abcam), rabbit monoclonal anti-GLUT3 (1:100, Abcam), or rabbit monoclonal anti-GLUT4 (ab313775, 1:100, Abcam) overnight at 4° C. To visualize antibody binding, sections were incubated with goat anti-rabbit IgG Alexa Fluor 647 secondary antibody (A-21245, 1:750, ThermoFisher) for one hr. To quench autofluorescence from lipofuscin and non-lipofuscin sources, sections were also treated with TrueBlack® (92401, Cell Signaling Technology) and TrueVIEW (SP-8400-15, Vector), respectively, as described by the manufacturers. Images of the labyrinth zone were acquired at 40x using a Leica TCS SP8 confocal microscope. Colocalization of GLUTs and WGA was evaluated using Manders’ Coefficient with Costes’ automatic threshold, calculated by ImageJ JACoP Software. Images from two locations (290 µm by 290 µm each) in the labyrinth zone were evaluated from each placenta.

### Placental Perfusion

*Ex vivo* placental perfusion (D’Errico et al. 2019) was performed on placentas from a subset of dams (n=8) to measure the real-time transport of glucose across the placenta. The gravid left horn was isolated and placed in low glucose (60 mg/dL) physiological salt solution (LG-PSS: 119 mM NaCl, 4.7 mM KCl, 1.17 mM MgSO_4_·7H_2_O, 1.6 mM CaCl_2_·2H_2_O, 1.18 mM NaH_2_PO4, 24 mM NaHCO_3_, 3.3 mM glucose, and 0.03 mM EGTA) (pH 7.4) under a dissection microscope. A uterine-placental-fetal unit was selected from the center of the uterine horn based on the length of the uterine artery and separated from the rest of the horn. The amniotic membrane was removed from the fetal surface revealing the umbilical vein and artery. The fetus was excised, and the maternal-placental unit was placed in a microvessel chamber (Living Systems Instrumentation, Fairfax, VT) filled with cold (4°C) LG-PSS. The distal and proximal ends of the uterine artery were cannulated and secured to glass cannula (175-200 µm) with nylon sutures. The umbilical vein and artery were cannulated and secured to blunt stainless steel needles (25G) with silk sutures. The microvessel chamber was warmed to 37 °C and the placenta was perfused with LG-PSS by a gravity-fed system set at 80 mmHg for the proximal uterine artery and 50 mmHg for the umbilical artery. Following 20 min equilibration and effluent collection, uterine inflow was increased to 200 mg/dL glucose (119 mM NaCl, 4.7 mM KCl, 1.17 mM MgSO_4_·7H_2_O, 1.6 mM CaCl_2_·2H_2_O, 1.18 mM NaH_2_PO4, 24 mM NaHCO_3_, 11 mM glucose, and 0.03 mM EGTA). Umbilical vein effluent was sampled, weighed, and glucose concentration evaluated using a glucometer every 10 min for 3 hr. Glucose flux (μmol/min) across the placenta was calculated and normalized to placental weight using the equation below:

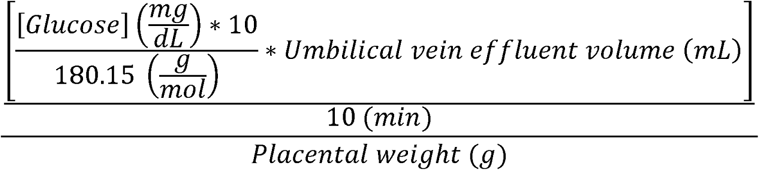

### Statistical Analyses

Statistical analyses were performed and graphs generated using GraphPad Prism v10 (Boston, MA). Maternal characteristics and litter size were analyzed using unpaired t-tests. Other litter characteristics and GLUT analyses are represented as a litter average for each dam. Data stratified by sex was analyzed by two-way ANOVA with Tukey’s post-hoc analysis. Fetal glucose levels below the limit of detection (20 mg/dL) were imputed as LOD/2 (Cary et al. 2023; Hornung and Reed 1990). Placental perfusion assays were analyzed by area under curve. Data are presented as mean ± SEM and deemed significant if p≤0.05. Outliers were manually removed from the data set and identified as values that were beyond two standard deviations of the mean.

## Results

### Maternal and Litter Characteristics

Maternal and litter characteristics were recorded on GD 20. Exposure to nano-TiO_2_ had no significant effects on maternal weight, mean arterial blood pressure, litter size, or fetal weight (Table 1). We have previously demonstrated a significant decrease in non-fasted fetal weights after exposure to nano-TiO_2_ in our larger project cohort of n=23–26 litters: 4.21 ± 0.08 g in exposed group compared to 4.39 ± 0.04 g in unexposed controls group. However, fetal weight was not significantly different between groups in this subset cohort (Table 1). Interestingly, placental weight was significantly increased in the exposed group compared to controls (Figure 2A), yet placental efficiency remained unaffected (Figure 2B). Taken together, gestational inhalation of nano-TiO_2_ impacts gross placental morphology.

**Figure 2.**
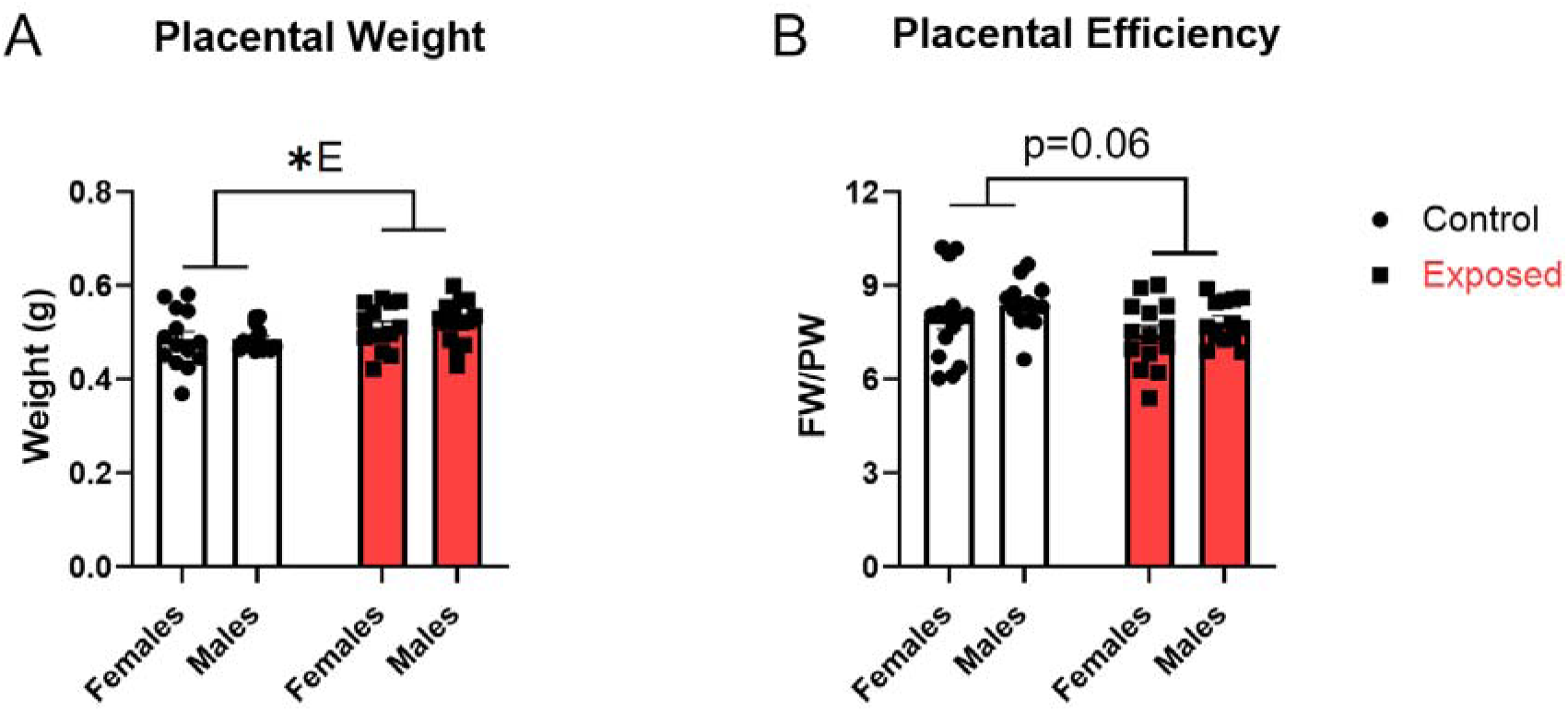
Effects of nano-TiO_2_ inhalation on placental morphometric parameters. Placental weight (PW) and placental efficiency were measured on GD 20. Placental efficiency was calculated by dividing average fetal weight (FW) by average PW. Data are mean ± SEM, n=14-15 litters/treatment group. ^*E^Significant exposure factor (p≤0.05) as determined by two-way ANOVA.

**Table 1.** Effects of nano-TiO_2_ inhalation on maternal and litter characteristics. Maternal weight, mean arterial pressure (MAP), litter size, and fetal weight were recorded on GD 20. Data are mean ± SEM, n=14-26 dams or litters/treatment group.

|  | **Control** |  | **Exposed** |  |
| --- | --- | --- | --- | --- |
| Maternal weight (g) (n=26) | 299.4 ± 4.9 | | 295.6 ± 3.8 | |
| MAP (mmHg) (n=9-11) | 75.8 ± 1.8 | | 78.5 ± 3.3 | |
| Litter size (n=19-26) | 10.2 ± 0.4 | | 11.1 ± 0.3 | |
|  | **Females** | **Males** | **Females** | **Males** |
| Fetal weight (g) (n=14-15) | 3.9 ± 0.1 | 4.0 ± 0.1 | 3.8 ± 0.1 | 3.9 ± 0.1 |

### Circulating Glucose and Insulin

Maternal and fetal blood glucose and maternal insulin were evaluated on GD 20. Maternal blood glucose was unaffected by gestational exposure to nano-TiO_2_ (Figure 3A). Interestingly, we observed a trending increase in fetal blood glucose (p=0.08) in exposed litters compared to control (Figure 3B). Additionally, maternal plasma insulin was not altered by exposure (Figure 3C). Overall, gestational inhalation of nano-TiO_2_ did not have a significant effect on systemic metabolic markers.

**Figure 3.**
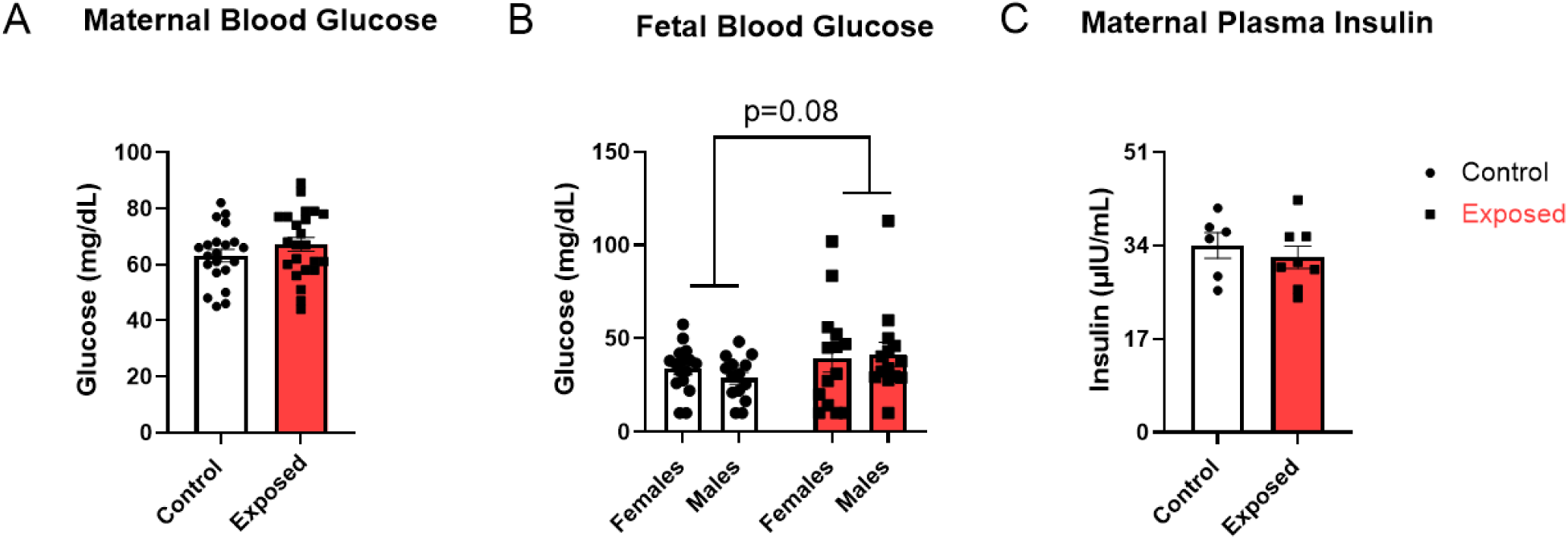
Effects of nano-TiO_2_ inhalation on systemic metabolic markers. Maternal (A) and fetal blood glucose (B) were measured using a glucometer and maternal plasma insulin (C) was evaluated using an immunoassay on GD 20. Data are mean ± SEM, n=8-23 dams or litters/treatment group. Analyzed two-way ANOVA.

### *GLUT* Expression

Placental mRNA expression of *Glut1* was unaffected by gestational exposure to nano-TiO_2_ (Figure 4A). Conversely, *Glut3* had significantly lower expression in exposed placentas compared to control (Figure 4B). There were no sex-related changes in *Glut3* expression. Exposure to nano-TiO_2_ demonstrated a sex-related effect on *Glut4* expression, indicating that placental *Glut4* mRNA decreased in females and increased in males. (Figure 4C). Collectively, these results demonstrate that gestational inhalation of nano-TiO_2_ differentially impacts glucose transporter expression at the transcriptional level, with sex-related regulation of *Glut4*.

**Figure 4.**
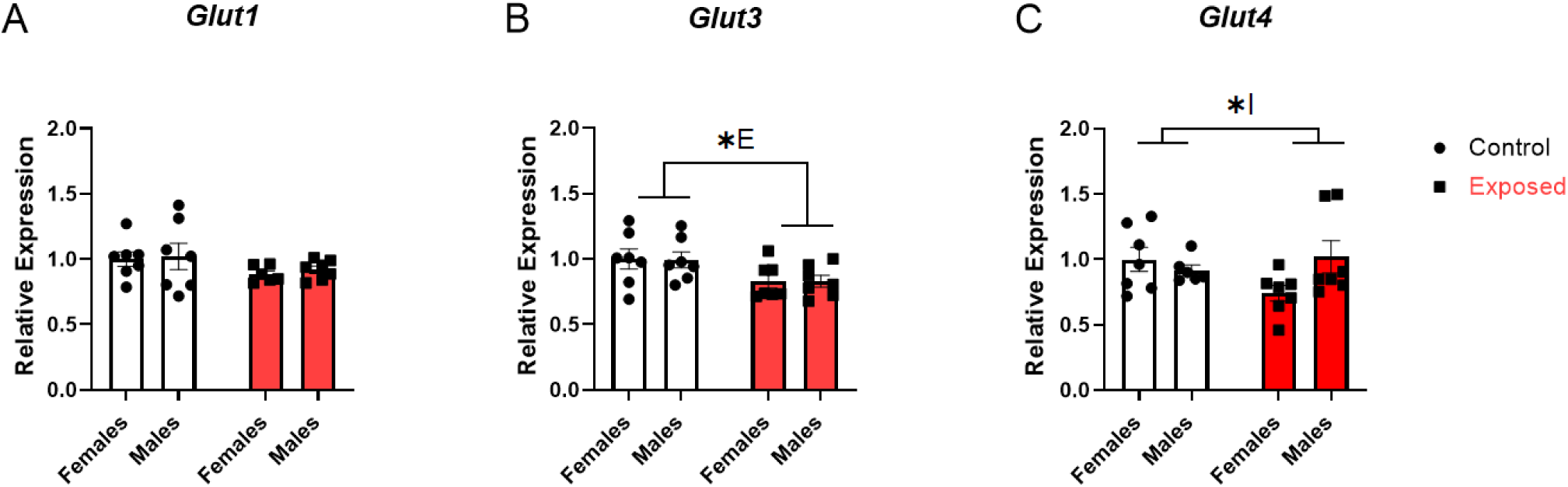
Effects of nano-TiO_2_ inhalation on *Glut* gene expression. GD 20 placentas were processed for RNA isolation and *Glut 1* (A), *3* (B), and *4* (C) gene expression was evaluated using RT-qPCR with *B-actin* as a housekeeping gene. Data are mean ± SEM, n=6-7 litters/treatment group. ^*E^Significant exposure factor, ^*I^Significant interaction factor (p≤0.05) as determined by two-way

Immunohistochemistry visualized the abundance of GLUT protein in the labyrinth zone. GLUT1 protein was significantly lower in exposed placentas as compared to control, as evidenced by a significant reduction in DAB staining (9% decrease) (Figure 5A and B). GLUT3 protein expression was unaffected by exposure. A significant increase in GLUT4 protein expression was identified as shown by an increase in DAB staining (Figure 5A and B). Post-hoc analyses revealed a more pronounced effect of nano-TiO_2_ on GLUT4 in female placentas (91% increase) compared to males (61% increase) (Figure 5B). Overall, these data suggest that gestational inhalation of nano-TiO_2_ negatively affects the expression of GLUT1, but a biological compensation occurs with an increase in overall protein expression of GLUT4.

**Figure 5.**
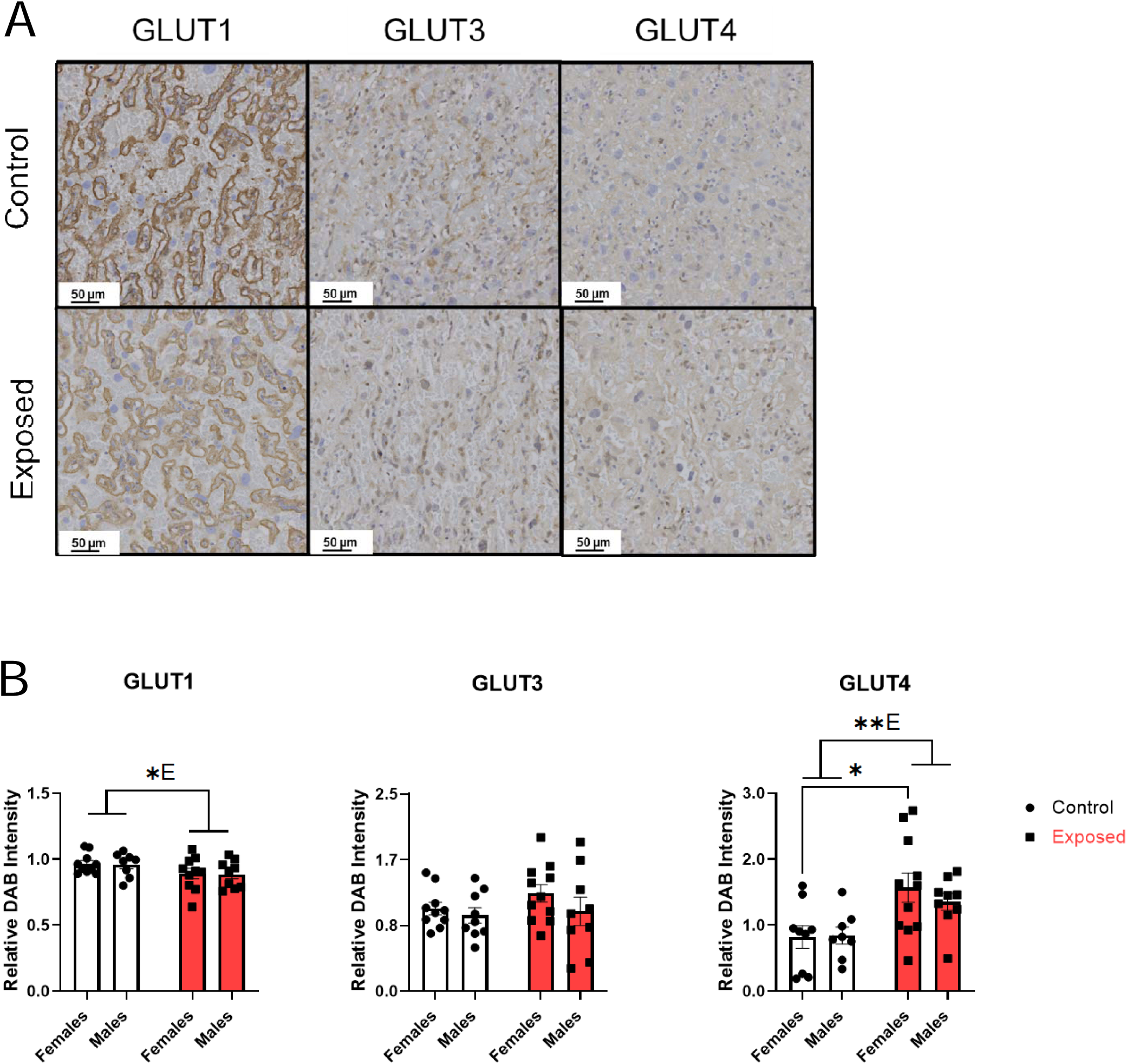
Effects of nano-TiO_2_ inhalation on placental GLUT protein expression. GD 20 placentas were histologically stained for GLUT1, 3, and 4. DAB chromogen substrate was used to visualize a positive signal for the protein and representative images were selected (A). DAB intensity was measured using ImageJ Fiji (B). Data are mean ± SEM, n=9-11 litters/treatment group. ^*E^Significant exposure factor (p≤0.05), **E (p<0.01); *p≤0.05 as determined by two-way ANOVA and Tukey’s post-hoc analysis.

Immunofluorescence/confocal microscopy was performed to evaluate membrane localization of GLUTs. Membrane localization of GLUT1 (Figure 6A) in the placenta was significantly increased with exposure to nano-TiO_2_ compared to control. Specifically, female placentas demonstrated a more marked increase in GLUT1 membrane localization (31% increase) compared to males (17% increase). GLUT3 membrane localization was unaffected by gestational exposure to nano-TiO_2_ and there were no sex-related differences in baseline membrane localization (Figure 6B). GLUT4 membrane localization was also increased in response to nano-TiO_2_ but there were no sex-related effects (Figure 6C). Taken together, gestational inhalation of nano-TiO_2_ enhances the membrane localization of glucose transporters, indicating an attempt to increase glucose uptake in the labyrinth zone of the placenta, allowing for transport to the fetus.

**Figure 6.**
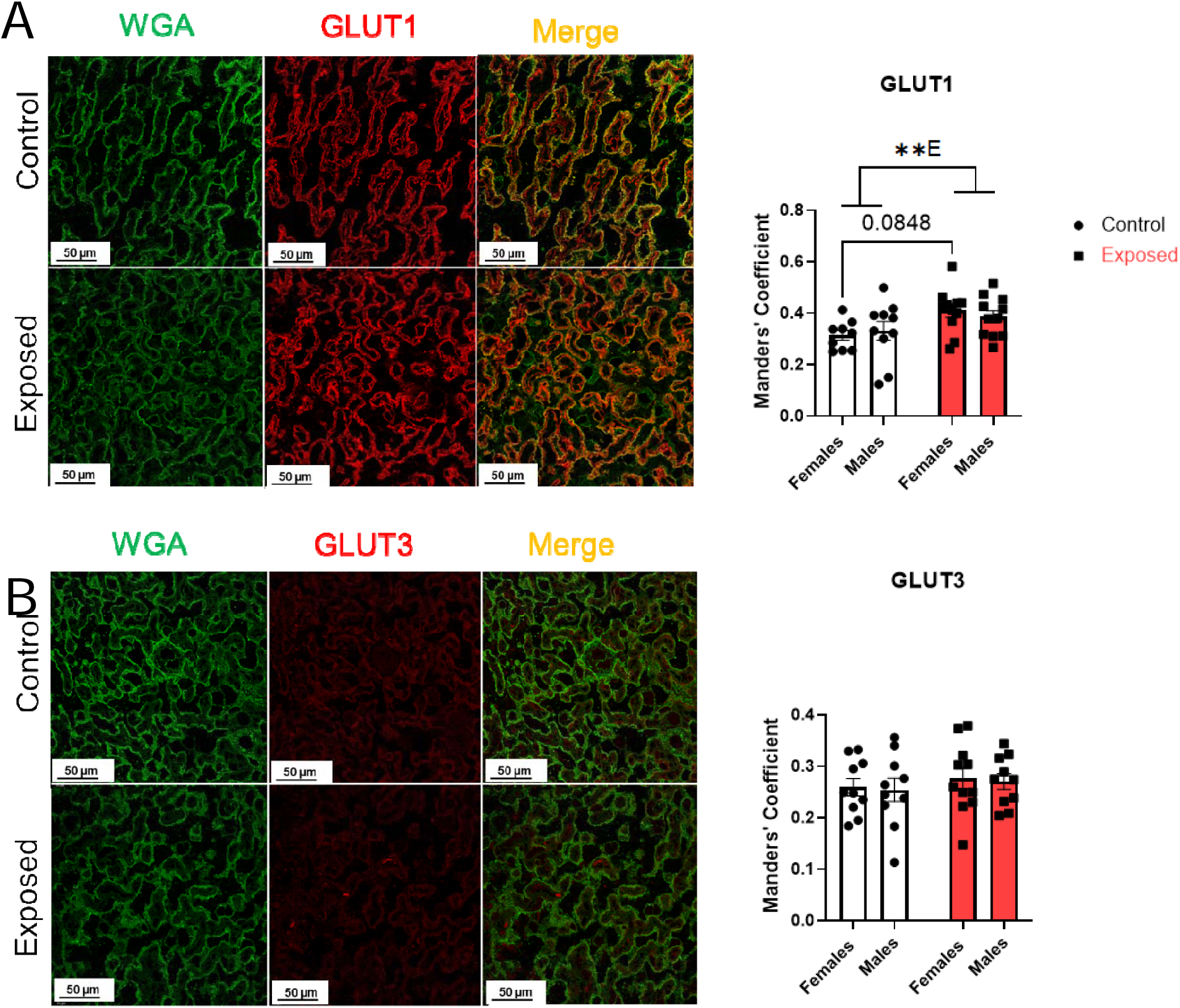
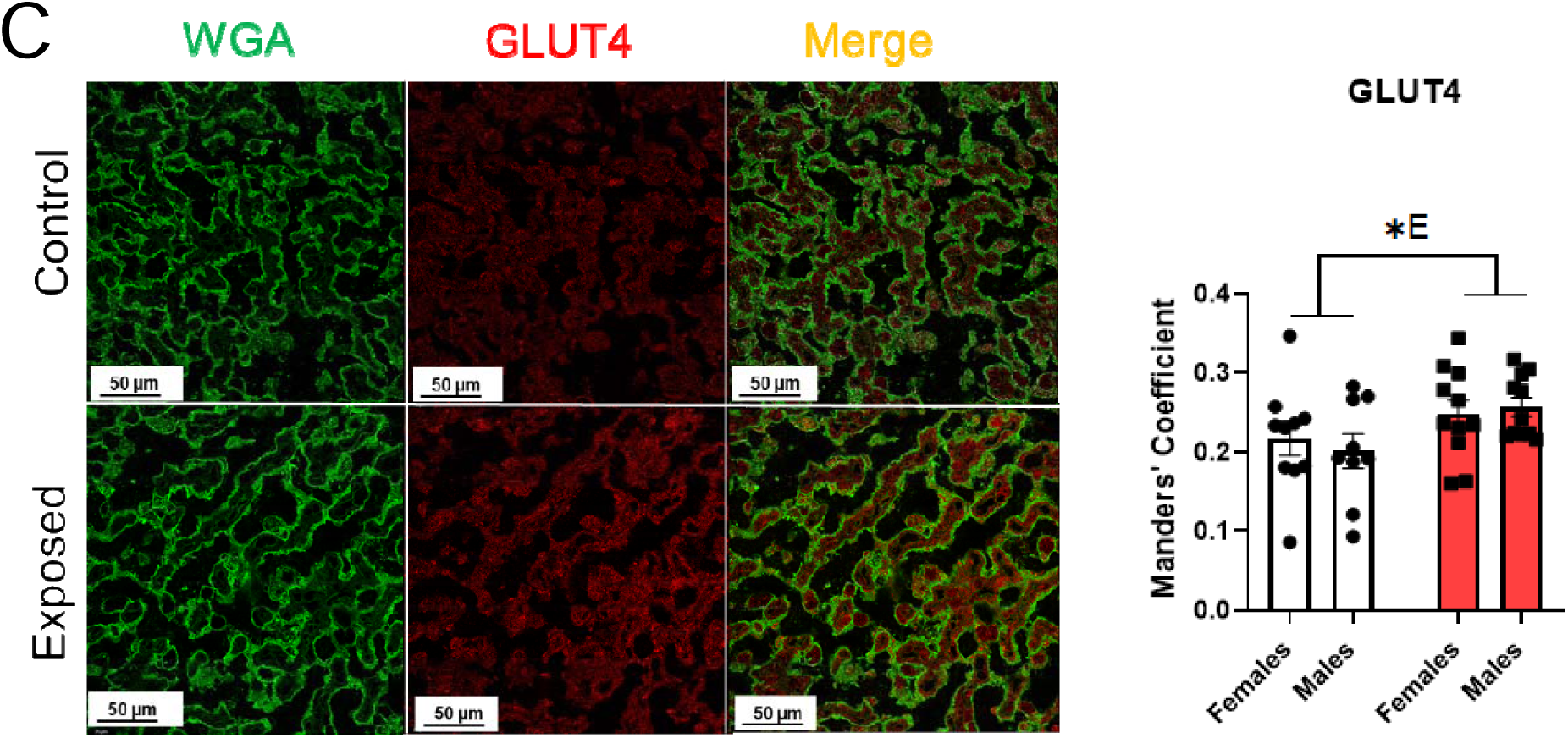
Effects of nano-TiO_2_ inhalation on placental GLUT membrane localization. GD 20 placentas were histologically stained with membrane marker, Wheat germ agglutinin (WGA), GLUT1 (A), 3 (B), and 4 (C). Alexa Fluor (AF) 488 was used to visualize WGA (green), AF647 used to visualize GLUTs (red), and representative images were collected. Manders’ Coefficient was calculated to quantify the co-localization of each transporter with WGA using ImageJ JaCoP software. Data are mean ± SEM, n=9-11 litters/treatment group. ^*E^Significant exposure factor (p≤0.05), **E (p<0.01) as determined by two-way ANOVA and Tukey’s post-hoc analysis.

### Placental Perfusion

To further investigate glucose transport capacity, glucose flux was evaluated via placental perfusion (Figure 7A). Placentas exposed to nano-TiO_2_ were associated with a significantly higher glucose flux compared to naïve control (AUC 95% CI: 77.4 to 127.5 vs. 39.1 to 73.6, respectively) (Figure 7B). This data corroborates our observations in GLUT localization, demonstrating that these molecular changes have functional consequences. Furthermore, gestational inhalation of nano-TiO_2_ leads to increased glucose transport across the placenta, into the fetal compartment.

**Figure 7.**
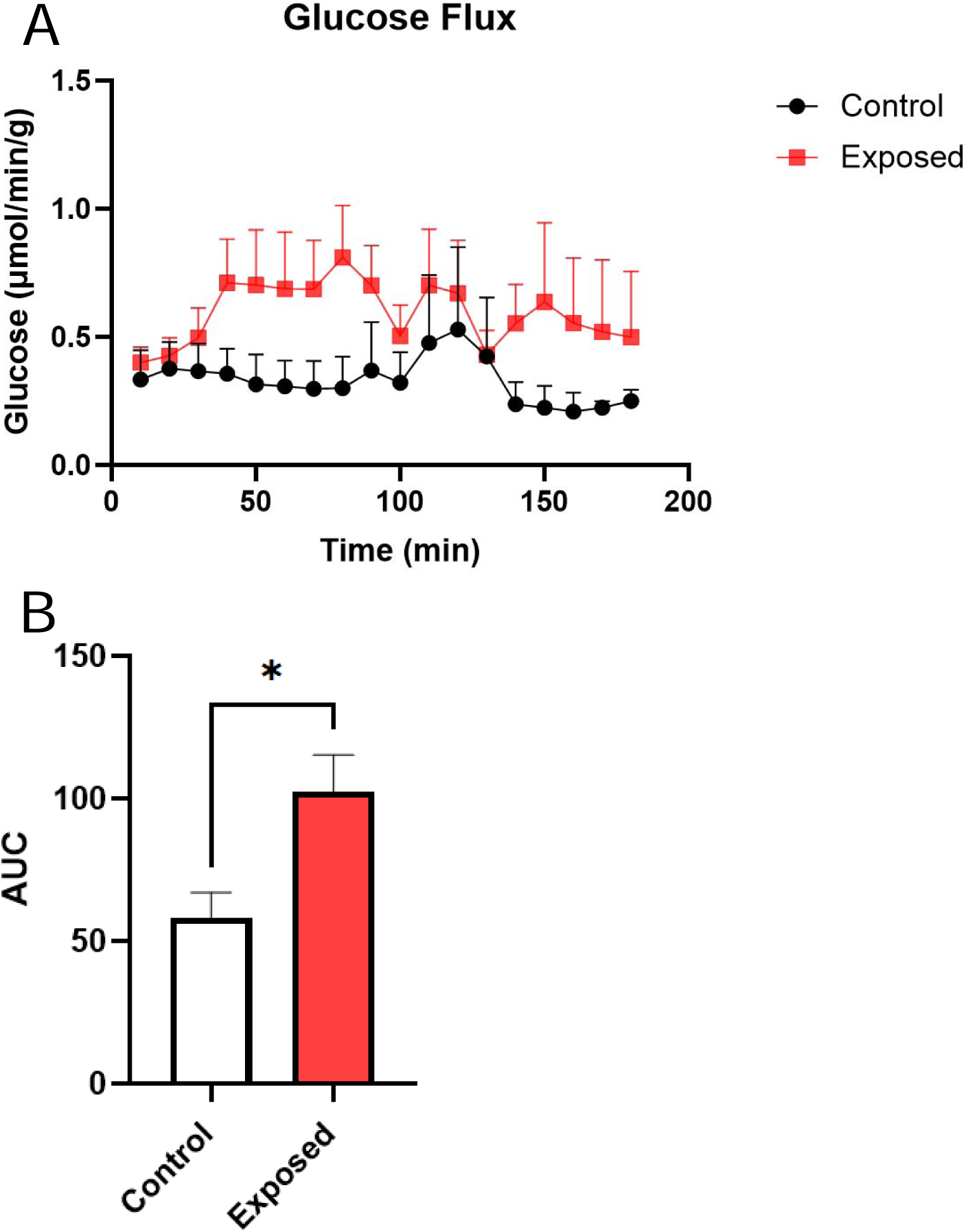
Effects of nano-TiO_2_ inhalation on placental glucose flux. GD 20 placentas were perfused through the uterine artery and umbilical artery with 60 mg/dL glucose. Following equilibration, uterine inflow was increased to 200 mg/dL. Umbilical vein effluent was sampled every 10 min for 3 hr to calculate glucose flux (A). Data are mean ± SEM, n=4-5 placentas/treatment group, analyzed by area under curve (AUC) (B).

## Discussion

Although environmental exposures to toxicants during pregnancy are frequently linked to adverse fetal outcomes, these associations often overlook the robust adaptive capacity of the placenta. As the maternal-fetal interface, the placenta can sense and respond to hostile conditions in ways that may partially preserve fetal development. The objective of this study was to assess placental glucose transport capacity after inhalation of ultrafine particulate matter during pregnancy to determine the mechanisms underlying the development of FGR. Contrary to our hypothesis that exposure would result in decreased glucose transport, molecular and histological changes in glucose transport mechanisms resulted in increased glucose flux across the placenta and into the fetal compartment.

Placental transport of glucose is gradient-dependent, resulting in increased glucose flux given high nutrient availability in the maternal compartment (Freinkel 1980). For this reason, the fetal hyperglycemia attributed to gestational diabetes is often associated with fetal overnutrition and overgrowth (Oken and Gillman 2003). In our model, we see increased placental glucose flux in the absence of gestational diabetes or fetal overgrowth. Under our current exposure conditions, we observed reduced *Glut3* mRNA and total GLUT1 protein expression, suggesting that inhalation of nano-TiO_2_ aerosols throughout gestation has an adverse effect on placental glucose transport. Prior *in vivo* studies with fine particulate matter exposure have also reported decreases in GLUT1 expression and reduced GLUT1 membrane localization (Villarroel et al. 2025), whereas we observed increased GLUT1 localization, which may be reflective of their pre-gestation exposure paradigm. Further analysis of transporter localization reveal that GLUT4 plays a major role in the resulting adaptations we reported, showing increases in both protein and membrane localization, a process known to be regulated by insulin signaling (Leto and Saltiel 2012). Although maternal insulin levels were unaffected, it is possible that fetal insulin contributed to these changes; however, not assessed in the present study. Exposure to fine particulate matter in rodent models has also been associated with increased GLUT3 expression, with no implications for GLUT localization (Ganguly et al. 2024; Zhu et al. 2021). These studies utilize samples of varying particle sizes and chemical composition, indicating that alterations in GLUT abundance may also depend on the physiochemical properties of ambient air pollutants.

It is important to note that many air pollution studies rely on inherently heterogeneous ambient air samples. These samples often contain toxicants from diverse sources including: heavy metals, chlorinated pesticides, gaseous components, and other contaminants (Ganguly et al. 2024). As it is difficult to stratify the toxicological outcomes, these particulate chemistries and physicochemical properties may cloud the mechanism of toxicity. Our focus narrows these variables to elucidate the relationship between maternal pulmonary xenobiotic particle exposure and the development of FGR. In our study, the use of nano-TiO_2_ allows us to recapitulate a narrow size fraction of ultrafine particulate matter in the ambient air.

The mechanisms for the development FGR in our nanoparticle model remain unclear; however, given the reduced uteroplacental perfusion previously demonstrated, it is plausible that impaired oxygen availability is a contributing factor (Fournier et al. 2019; Stapleton et al. 2013). Low birth weight often reflects reduced uteroplacental blood flow (Dumolt et al. 2021; Wang 2010), leading to inadequate oxygen delivery to the fetus and mitochondrial dysfunction in the developing pancreas (Simmons 2012). Consequently, low birth weight is linked to metabolic syndrome in adulthood, including the risk for insulin resistance, obesity, and type 2 diabetes (Barker et al. 1997; Fall et al. 1995; Law et al. 1992; Liao et al. 2020; Loos et al. 2001; Rich-Edwards et al. 1999). We have also shown that the placenta exhibits mitochondrial dysfunction and a metabolic shift away from oxidative metabolism to glycolytic metabolism (Seymore et al. 2025), a hallmark response to hypoxia (Illsley et al. 2010; Innocenti et al. 2024; Kierans and Taylor 2021). Because this metabolic shift requires increased glucose consumption by the placenta to maintain ATP concentrations, it requires enhanced GLUT expression and activity to support enhanced glucose uptake (Kierans and Taylor 2021). This adaptive response may inadvertently lead to excess glucose uptake and transport, an overcompensation that results in elevated fetal glucose levels. Lastly, we have previously identified the translocation of inhaled nano-TiO_2_ to the placenta in our FGR model (D’Errico et al. 2022), therefore, we cannot rule out that our results may be associated with the translocation of ultrafine particulates i.e., nanoparticles (<100 nm in diameter), from the air-blood barrier (Hussain et al. 2011), thus, eliciting direct particle effects systemically within maternal physiology (i.e., neurological or endocrine adaptations) or locally on the uterine vasculature or placental morphology (Stapleton et al. 2015a).

Despite the strengths of this study, including the use of *ex vivo* placental perfusion for real-time assessment of glucose transport, a few limitations should be noted. First, unlike most of the datasets presented, our placental perfusion data were not stratified by fetal sex. As we did not identify sex-related changes in GLUT membrane localization, we did not expect to observe sex-related variations in glucose flux across the placenta. Additionally, there are anatomical differences that may limit the translation of our outcomes. The human placental barrier contains one syncytiotrophoblasts layer, whereas the rat placental barrier consists of two layers (Furukawa et al. 2018). In this structure, GLUT1 is predominantly localized to the apical side of the maternal-facing layer and the basolateral side of the fetal-facing layer, GLUT3 is mainly found on both the apical and basolateral sides of the maternal facing layer, and GLUT 4 is mainly intracellular (Shin et al. 1997; Takata and Hirano 1997). Knowing the specific translocation pathways would offer more insight into whether these changes mostly benefit cellular uptake and metabolism or cellular uptake and export to the fetus. Accordingly, the polarization of these transporters should be considered in future studies.

Overall, we have demonstrated that the inhalation of nanoparticles alters placental glucose transport in a way that increases fetal glucose bioavailability. These changes can predispose the fetus for metabolic reprogramming, underscoring the potential for long-term environmental-induced metabolic risks. By focusing on ultrafine particulate exposure and using *ex vivo* placental perfusion, this work identifies transporter-specific adaptations in the placenta that provide insight into fetal responses to environmental pollutants. Furthermore, our findings expand the understanding of how inhaled nanoparticles affect placental nutrient transport and point to potential molecular targets for mitigating adverse fetal developmental outcomes.

## Author contributions: CRediT

**Talia Seymore**: Writing – original draft, Visualization, Validation, Project administration, Methodology, Investigation, Formal analysis, Data curation, Conceptualization. **Sara Hoffmann**: Investigation, Formal analysis, Writing – Review & Editing. **Pedro Louro:** Validation, Investigation. **Carol Gardner:** Methodology, Validation, Resources, Writing – Review & Editing, Supervision. **Michael Goedken:** Investigation, Resources, Writing – Review & Editing. **Phoebe Stapleton:** Writing – Review & Editing, Supervision, Resources, Project administration, Methodology, Funding acquisition, Conceptualization, Validation.

## Acknowledgments

This work was supported by the National Institutes of Health [NIH R01ES031285, T32ES007148, and P30 ES005022]. The authors also acknowledge the contributions of lab managers Tanisha Brunson-Malone and Marianne Polunas, and PhD student Samantha Adams for performing the animal exposures.

