## Supplemental Figure 1 for "Inhalation of nanoparticles during pregnancy enhances placental glucose transport in rats"

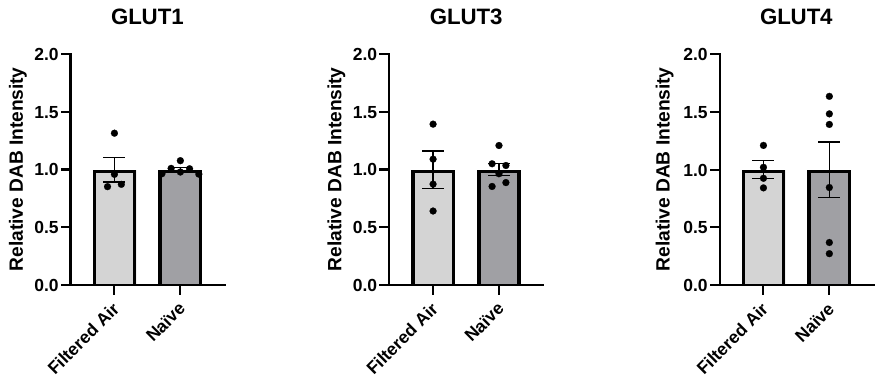

Supplemental Figure 1. GLUT protein expression on naïve placentas and those exposed to filtered air. GD 20 placentas were histologically stained for GLUT1, 3, and 4. DAB chromogen substrate was used to visualize a positive signal for the protein. DAB intensity was measured using ImageJ Fiji as a semi-quantitative analysis of protein expression. Data are mean ± SEM, n=4-6 litters/treatment group, analyzed by unpaired t-test.
